# Significant shared heritability underlies suicide attempt and clinically predicted probability of attempting suicide

**DOI:** 10.1101/266411

**Authors:** Douglas M. Ruderfer, Colin G. Walsh, Matthew W. Aguirre, Yosuke Tanigawa, Jessica D. Ribeiro, Joseph C. Franklin, Manuel A. Rivas

## Abstract

Suicide accounts for nearly 800,000 deaths per year worldwide with rates of both deaths and attempts rising. Family studies have estimated substantial heritability of suicidal behavior; however, collecting the sample sizes necessary for successful genetic studies has remained a challenge. We utilized two different approaches in independent datasets to characterize the contribution of common genetic variation to suicide attempt. The first is a patient reported suicide attempt phenotype from genotyped samples in the UK Biobank (337,199 participants, 2,433 cases). The second leveraged electronic health record (EHR) data from the Vanderbilt University Medical Center (VUMC, 2.8 million patients, 3,250 cases) and machine learning to derive probabilities of attempting suicide in 24,546 genotyped patients. We identified significant and comparable heritability estimates of suicide attempt from both the patient reported phenotype in the UK Biobank (h^2^_SNP_ = 0.035, p = 7.12×10^−4^) and the clinically predicted phenotype from VUMC (h^2^_SNP_ = 0.046, p = 1.51×10^−2^). A significant genetic overlap was demonstrated between the two measures of suicide attempt in these independent samples through polygenic risk score analysis (t = 4.02, p = 5.75×10^−5^) and genetic correlation (rg = 1.073, SE = 0.36, p = 0.003). Finally, we show significant but incomplete genetic correlation of suicide attempt with insomnia (rg = 0.34 - 0.81) as well as several psychiatric disorders (rg = 0.26 - 0.79). This work demonstrates the contribution of common genetic variation to suicide attempt. It points to a genetic underpinning to clinically predicted risk of attempting suicide that is similar to the genetic profile from a patient reported outcome. Lastly, it presents an approach for using EHR data and clinical prediction to generate quantitative measures from binary phenotypes that improved power for our genetic study.

## Introduction

Suicide accounts for over 40,000 deaths a year in the United States alone and close to 800,000 deaths worldwide^1,2^. Suicide attempt and ideation affect a much larger proportion of the population with estimates of attempts 10-25 times the number of individuals that die from suicide (1 million in the US, 20 million worldwide^3^) and a 6-14% lifetime prevalence of suicidal ideation^4^. Despite preventative public health efforts, rates of suicide and suicidal behavior are increasing in the U.S., particularly among young adults^5^. Epidemiological and family studies imply a substantial genetic component with estimates of the heritability of suicide attempt as high as 55%^6–9^. However, large-scale genetic studies remain difficult due to challenges in phenotypic ascertainment and collecting large enough samples to have the power to identify replicable genetic associations or to directly estimate the proportion of heritability contributed from common genetic variation^10–13^. Thus, despite both the major public health impact and the strong evidence of heritability, the genetic architecture of suicide and suicidal behaviors remains poorly understood^14^.

The emergence of large-scale, population-based samples where participants are phenotypically screened and genetically interrogated provides opportunites to study the genetics of phenotypes at scale. While the vast majority of individuals who attempt suicide have been diagnosed with a psychiatric disorder^15,16^, the outcome is not limited to any single diagnosis. Previous work has pointed to genetic factors independent of diagnosis^17,18^, which make population samples particularly valuable for the study of suicide. The UK Biobank, has enrolled 500,000 individuals with extensive phenotypic and genetic data, including an online mental health assessment taken by over 157,000 participants. In this assessment, participants reported self-harm behaviors and specifically whether they have ever attempted suicide with the intent to end their lives. Among those questioned, over 3,000 participants responded “yes,” providing a large set of suicide attempt cases with genetic data currently available and a corresponding set of population matched controls.

Parallel efforts have been utilizing large-scale clinical data (diagnoses, medications, procedures, utilization, demographics, etc.) from electronic health records (EHR) to identify features associated with suicide attempt and to apply predictive analytics to assess risk of future suicidal behaviors^19–21^. The most recent efforts in this domain have applied machine learning with high accuracy (c-statistics above 0.8-0.9) and precision (above 0.8) for suicide attempts^20^ and death^19,21–23^. While the goal is often to predict a binary outcome (e.g., suicide attempt or death) an important product generated from these approaches is a posterior probability associated with the likelihood of the outcome occurring (e.g., probability of attempting suicide at any point in time). These probabilities can be generated for every patient with relevant data at hospital- or system-scale, regardless of whether they have the outcome or not, and are well-suited to serve as quantitative phenotypes for genetic studies. Integrating predictive analytics of suicide from EHR data and genetic data allows for an opportunity to provide meaningful quantitative phenotypes for all genotyped patients and not rely on the small subset of patients who have already engaged in suicidal behavior.

In this work, we exploit both the large-scale population genetic sample from the UK Biobank and a hospital based EHR and genetic sample from the Vanderbilt University Medical Center (VUMC) to study the genetics of patient reported suicide attempt along with clinically predicted probability of suicide attempt derived from validated algorithms of suicide risk^20^. We perform genome-wide association analyses on both samples, estimate heritability of each and calculate the genetic correlation between them and across hundreds of other traits to interrogate the genetic contribution to suicide attempt and predicted risk of attempting suicide. These analyses directly address how common genetic variation contributes to suicide attempt, whether a biological basis underlies clinical predictions of suicide attempt and whether clinical prediction can be used to increase the power of genetic studies by adding a quantitative dimension to a dichotomous phenotype. Importantly, the approach is generalizable and may be applied equally well to a wide variety of medical diagnoses or traits.

## Methods

### Genotyping and quality control of the UK Biobank sample

Genotyping and imputation procedures for the UK Biobank dataset have been previously described^24^. Briefly, two genotyping arrays, the UK Biobank Axiom Array (n=438,427) and the UK BiLEVE Axiom Array (n=49,950), were used to create the final genotype release of 805,426 loci for 488,377 individuals. Genotype quality control was performed before the data were released publicly, including removing participants with excess heterozygosity or missingness rate, and removing markers showing effects related to batch, plate, sex or array, or those demonstrating discordance across control replicates. Imputation was performed using a reference panel derived from the Haplotype Reference Consortium (HRC), the UK10K and 1000 Genomes datasets. Pre-phasing was leveraged to gain computational efficiency by imputing haploid genotypes for each sample. A total of 670,739 variants were used for pre-phasing and imputation if they were present on both arrays, passed genotype QC in all batches, had MAF > 0.0001, and had missingness < 5%. A total of 39,313,024 variants present in HRC were imputed.

Genome-wide association analysis was conducted using logistic regression with Plink v2.00a on the set of imputed variants from 337,199 unrelated individuals of white British ancestry in the UK Biobank. The following covariates were used for the analysis: age, sex, the first four genetic principal components, and array, which denotes whether an individual was genotyped with the UK Biobank Axiom Array or the UK BiLEVE Axiom Array. Variants present on only one array were run without array as a covariate. Phenotypes were defined using UK Biobank Data-Field 20483 (Ever attempted suicide). Cases are “yes” responses (n=2,433), and controls are either “no” responses, or any other response (N=334,766). Imputed dosages were filtered for having minor allele frequency greater than 1% and imputation INFO score > 0.3 resulting in a final set of 7,797,387 variants.

### Genotyping and quality control of the VUMC BioVU sample

The VUMC has a patient population of nearly 3 million individuals for whom clinical data are stored and managed in EHR. DNA has been collected on over 250,000 of these patients (as of February 2018), and linked to clinical data that are de-identified for use in genetic studies^25^. For this study, individuals had been previously genotyped on three different Illumina platforms and experiments were performed at different times. The samples consisted of 24,262 individuals genotyped on the Illumina MEGA^EX^ platform consisting of nearly 2 million markers, 6,483 individuals genotyped on the Illumina Omni1M array covering nearly 1 million markers, and 4,035 individuals were genotyped on the Illumina Human660W array covering 600,000 markers. We removed samples with greater than 2% missingness or abnormal heterozygosity (|Fhet| > 0.2). Variants were excluded if they had greater than 2% missingness or Hardy-Weinberg equilibrium p-value < 5×10^−5^. We excluded SNPs with minor allele frequency less than 1% and those not genotyped in HapMap2. Genotype imputation was performed using the prephasing/imputation stepwise approach implemented in IMPUTE2 / SHAPEIT using 1000 genomes phase I reference panel. Variants were excluded for having low imputation quality (INFO < 0.3). A set of SNPs QC-ed and pruned for linkage disequilibrium was used to calculate relatedness and principal components of ancestry. For pairs of highly related individuals (pihat > 0.2), one was randomly excluded. Ancestry components were used to define a homogenous population sample and included as covariates in association analysis to account for ancestry confounding. MEGA samples were genotyped in five batches and variants were removed if having significantly differing frequencies (p < 5×10^−5^) between any batch and the rest of the sample within homogenous sets of individuals. Individuals having been genotyped on multiple platforms were retained only in one with preference for being on the MEGA array.

### Predicted probability of attempting suicide and feature quantification

The EHR-based phenotyping of suicide attempt and machine learning derived-phenotyping algorithm used here were adapted from a published predictive model of suicide attempt risk using clinical EHR data at VUMC^20^. Briefly, clinical data were collected from the de-identified repository known as the VUMC Synthetic Derivative (SD)^25^. Candidate charts were identified using self-injury International Classification of Diseases, version 9 (ICD-9) codes (E95x.xx) for all adults in the SD. Cases of suicide attempts were identified through multi-expert chart review on a candidate list of 5,543 charts with self-injury codes to identify 3,250 adults (aged 18 or older) with expert-validated evidence of self-harm with suicidal intent. A cohort of 12,695 adults with a minimum of three visits to VUMC were drawn from the general population as the control comparison. Clinical data were preprocessed to support clinical prediction/phenotyping including demographics; clinical diagnoses grouped from individual ICD-9 codes to Center for Medicare and Medicaid Servicers Hierarchical Condition Categories (CMS-HCC); medications grouped to the Anatomic Therapeutic Classification, level V; healthcare utilization including counts of inpatient, outpatient, and emergency department visits for each year of the preceding five years^20^. Missing data were rare because the variables measured as counts – diagnoses, medications and visits – were imputed to zeroes if not present. Zip code used to calculate area deprivation index was missing in 6% of charts, body mass index was missing in 9.9%, race was missing in 3.6%, and date of birth used to calculate age was missing in 0.7%. Multiple imputation was used to impute missing values in those instances^26^.

As per our prior work^20^, Random forests have been shown to have superior discrimination performance in identifying suicide risk. With tuning parameters of 500 trees per forest and splits of the square root of the number of predictors at each node in the tree, the clinical phenotyping algorithm was trained via optimism adjustment with the bootstrap using 100 bootstraps^27^. The model used here differed from the published model only in that it did not censor clinical data n days (where n ranged from 7 to 730) preceding attempt. Therefore, discrimination performance was similar to the published models (AUC = 0.94 [0.93-0.95], sensitivity = 0.92, specificity = 0.82). The phenotyping algorithm was applied to 235,932 patients with genetic data in the biobank at VUMC (BioVU). Posterior probabilities were normalized using a rank-based inverse transformation and used as the quantitative phenotypes in genetic analyses (results remained stable when applying other normalization approaches, data not shown).

## Results

### Genome-wide association study (GWAS) of suicide attempt in UK Biobank

A total of 157,366 participants provided responses to an online mental health questionnaire as a follow up to initial phenotyping in the UK Biobank sample. Of these, 6,872 were asked this question from Data-Field 20483, Category: Self-harm behaviors, “Have you harmed yourself with the intention of ending your life?” Most participants were not asked this question as it required a positive response to a previous self-harm question. In total, 3,563 of 6,872 respondents indicated “yes”, 3,089 responded “no” and 220 preferred not to answer. In an effort to maximize power and because the phenotype is rare, we included all UK Biobank participants as controls except for those responding yes to attempting suicide, this includes those that did not take the mental health assessment at all and those who preferred not to answer. After reducing our sample to a set of homogenous Caucasians, we retained case-control data of 2,433 individuals having attempted suicide and 334,766 controls (see Methods). Nearly 8 million imputed dosages based on the HRC reference panel were tested after filtering for INFO > 0.3 and MAF > 1%. No variants reached our genome-wide significance threshold of p < 5×10^−8^ (Figure 1a-b). SNP-based heritability was estimated by LD-score regression^28^ using the prevalence of suicide attempt of the participants taking the online questionnaire to convert to liability scale. We identified significant SNP-based heritability (h^2^_SNP_ = 0.035, SE = 0.01, p = 7.12×10^−4^, Table 1) in the patient reported suicide attempt phenotype.

**Table 1.** Results from heritability estimates using LD-score regression of predicted probability of attempting suicide within each genotyping array in BioVU (first three rows), all of BioVU (fourth row) and patient reported suicide attempt in UK Biobank (fifth row).

| Sample | Cohort | $\lambda$ | mean $\chi^2$ | $h^2$ | SE | Z | p |
| --- | --- | --- | --- | --- | --- | --- | --- |
| BioVU | MEGA | 1.020 | 1.018 | 0.043 | 0.026 | 1.678 | $9.33 \times 10^{-2}$ |
| | 660 | 1.017 | 1.017 | 0.218 | 0.151 | 1.445 | $1.48 \times 10^{-1}$ |
| | Omni1M | 0.999 | 0.996 | 0.148 | 0.109 | 1.359 | $1.74 \times 10^{-1}$ |
| *BioVU* | *All* | *1.029* | *1.029* | *0.046* | *0.019* | *2.431* | *$1.51 \times 10^{-2}$* |
| *UK Biobank* | *All* | *1.038* | *1.038* | *0.035* | *0.010* | *3.385* | *$7.12 \times 10^{-4}$* |

**Figure 1.**
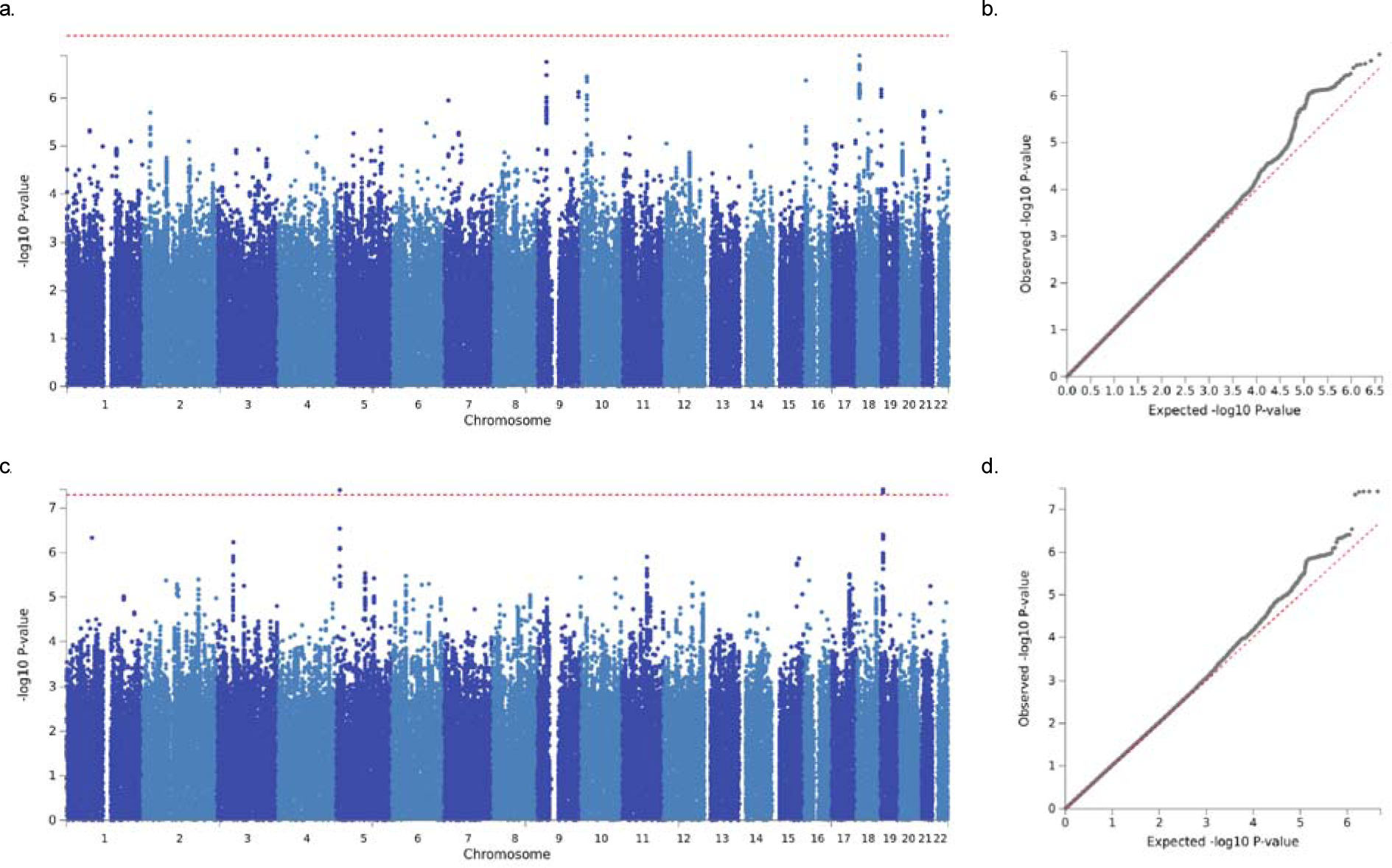
Genome-wide association results: a) Manhattan plot for UK Biobank participants attempting suicide vs all controls, red line represents p = 5×10^−8^. b) QQ-plot for UK Biobank suicide attempt. c) Manhattan plot for linear regression of predicted probability of attempting suicide in BioVU, red line represents p = 5×10^−8^. d) QQ-plot for predicted probability of attempting suicide in BioVU.

### Polygenic risk score analysis in clinically predicted risk of attempting suicide

After QC and filtering for a homogenous set of genotyped Caucasian patients from the biobank at VUMC (BioVU), we retained 24,546 patients with high quality genotyping data across three platforms of which 73 had attempted suicide based on expert chart review. We calculated polygenic risk scores using the effect sizes calculated from the UK Biobank GWAS across all SNPs that overlapped those genotyped on each of the platforms. Despite small numbers, we identified a significant increase of polygenic risk among patients with a chart validated suicide attempt compared to the rest of patients in BioVU in both the single largest dataset (n=18,128 patients, n=40 suicide attempts, p=0.016) and across the entire sample (n=24,546, n=73 suicide attempts, p=0.033). Leveraging EHR data on almost 3 million patients at VUMC, including 3,250 with chart validated suicide attempt, we adapted a previously published machine-learning based clinical prediction model^20^ (see Methods) to assign posterior probabilities of attempting suicide to all genotyped patients. We then tested the relationship between the UK Biobank based polygenic risk score and the predicted probability of attempting suicide using linear regression including four principal components and sex as covariates (including 20 principal components did not change results, data not shown). We identified a significant positive relationship between polygenic risk and predicted probability of attempting suicide in both the largest genotyped dataset (p = 9.96×10^−6^, t-stat = 4.42) and across all samples (p = 5.75×10^−5^, t-stat = 4.02), and all datasets showed positive direction (Table 2).

**Table 2.** Results from polygenic risk score analysis using UK Biobank GWAS summary statistics as discovery and testing aggregate genetic risk between BioVU patients having chart reviewed suicide attempt (left side) and quantitative probability of suicide attempt (right side) using logistic and linear regression, respectively. Columns are as follows; Est: regression estimate, SE: standard error, T: regression t-statistic, P: p-value.

| Sample | N | Validated attempt | Suicide attempt |  |  |  | Predicted risk of suicide attempt |  |  |  |
| --- | --- | --- | --- | --- | --- | --- | --- | --- | --- | --- |
|  |  |  | Est | SE | T | P | Est | SE | T | P |
| MEGA | 18,128 | 40 | 22.0 | 9.2 | 2.41 | 0.016 | 893.7 | 202.2 | 4.42 | $9.96 \times 10^{-6}$ |
| 660 | 2,965 | 17 | -16.2 | 31.4 | -0.52 | 0.607 | 491.7 | 424.5 | 1.16 | 0.247 |
| Omni1M | 3,453 | 16 | 33.9 | 24.0 | 1.41 | 0.157 | 44.2 | 375.9 | 0.12 | 0.906 |
| *Total* | *24,546* | *73* | *18.39* | *8.60* | *2.14* | *0.033* | *659.9* | *164.0* | *4.02* | *$5.75 \times 10^{-5}$* |

### GWAS of predicted probability of attempting suicide in BioVU

We next sought to identify specific variants contributing to predicted probability of suicide attempt in BioVU. Linear regression was performed on predicted probability of suicide attempt and allelic dosage including 4 principal components of ancestry for 9 million imputed variants after filtering out variants with minor allele frequency < 1% and imputation INFO scores < 0.3. Association was performed separately for each of the genotyping platforms, and inverse weighted meta-analysis was used to combine them with Plink^29^. We identified two genomic regions containing five variants surpassing genome-wide significance (p < 5×10^−8^) on chromosomes 5 and 19 with the most significant variants being rs12972617 and rs12972618 (p=3.81×10^−8^, beta=-0.063, Figure 1c-d). However, none of these variants replicated at nominal significance (p < 0.05) in the UK Biobank GWAS (Supplementary Table 1). We identified significant SNP based heritability (h^2^_SNP_ = 0.046, SE = 0.019, p = 0.015, Table 1) at around the same level as the patient reported outcome of suicide attempt used in the UK Biobank data.

### Genetic correlation of suicide attempt and other phenotypes

Both the GWAS of suicide attempt in UK Biobank and the GWAS of predicted probability of suicide attempt demonstrated significant heritability estimates of around 4%, and significant correlation was observed between predicted probability of suicide attempt and a polygenic risk score calculated from patient reported suicide attempt in UK Biobank. To further quantify the overlap between the genetic architecture of these two measures of the same trait we calculated genetic correlation and identified significant rg of 1.073 (SE = 0.36, z-score = 2.98, p = 0.002). We next assessed genetic correlation between GWAS summary statistics of our two suicide attempt phenotypes and 233 other GWAS traits^30^. Genetic correlations were performed for each phenotype separately and then meta-analyzed using Stouffer’s method of combining z-scores. In total, we performed 466 tests making our multiple test corrected significance threshold p < 1.07×10^−4^. Eight phenotypes surpassed this threshold in categories such as reproduction, sleep and psychiatric disorders (Figure 2). Specifically, we identified significant positive genetic correlation of suicide attempt with depressive symptoms^31^, neuroticism^31^, schizophrenia^32^, insomnia^33,34^, major depressive disorder^35^ and a combined phenotype of five psychiatric disorders^36^ as well as significant negative genetic correlation with age at first birth^37^ (Supplementary Table 2). Two traits showed nominally significant genetic correlation in both phenotypes but in opposite directions including intelligence (BioVU: rg = −0.53, p = 3×10^−4^, UK Biobank: rg = 0.19, p = 0.044) and years of schooling^38^ (BioVU: rg = −0.53, p = 3.3×10^−5^, UK Biobank: rg = 0.19, p = 0.007).

**Figure 2.**
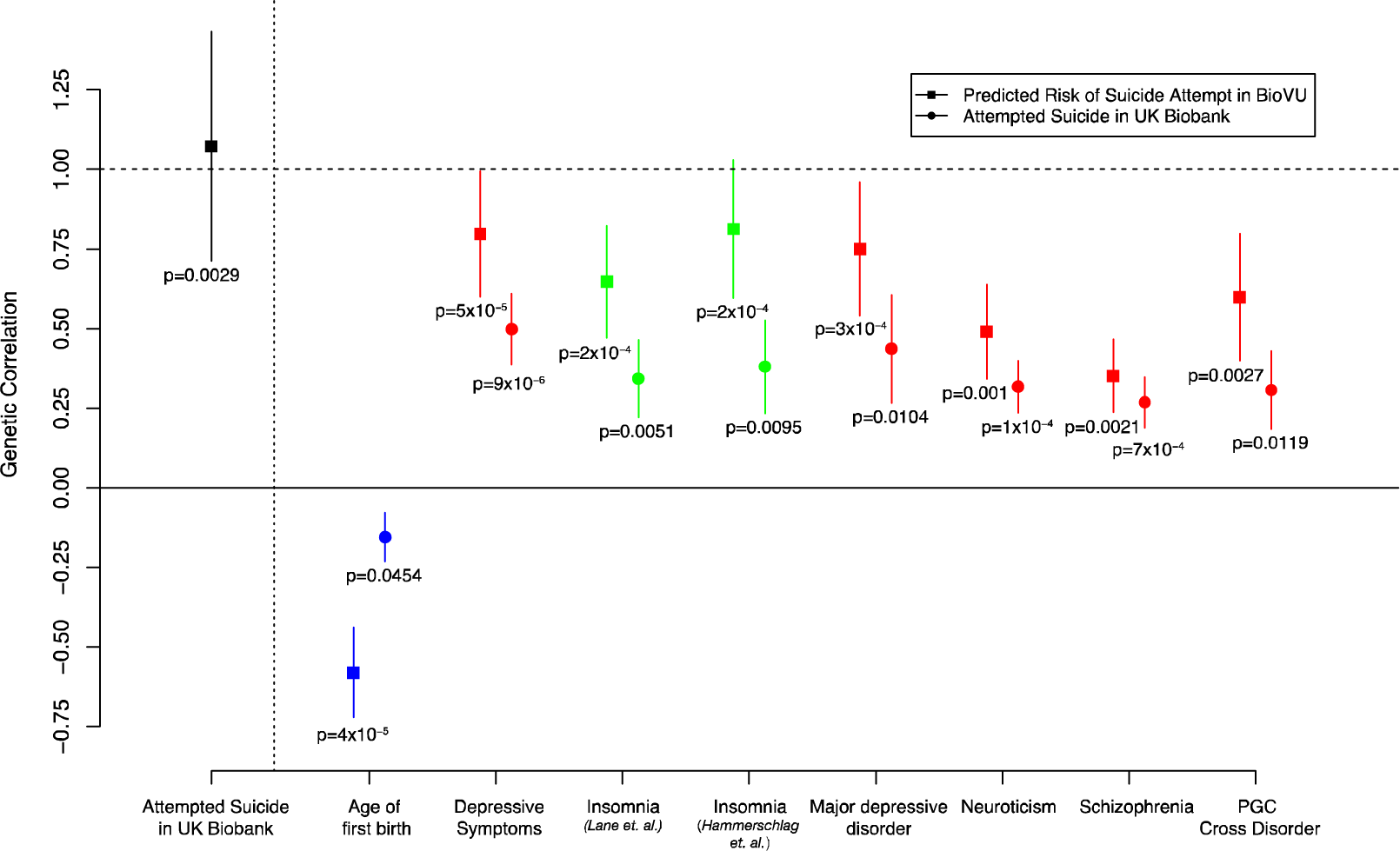
Genetic correlations and standard errors: Black point is rg between association of patient-reported suicide attempt in UK Biobank and predicted probability of attempting suicide in BioVU. Colored points represent set of phenotypes surpassing multiple test corrected significance of genetic correlation with suicide attempt after meta-analysis of UK Biobank and BioVU. Colors represent phenotype category (psychiatric = red, sleep = green, reproduction = blue). Square points are rg with predicted probability of suicide attempt in BioVU and circle points are rg with suicide attempt in UK Biobank. Complete rg results are in Supplementary Table 2.

## Discussion

We present two large-scale genetic analyses of suicide attempt based on population samples. One from a national effort with direct assessment of suicide attempt through online questionnaire and one from a hospital system where suicide risk was predicted based entirely on clinical features from EHR data. Both analyses demonstrated significant heritability estimates from common variation of around 4% and significant genetic correlation between them and with previously implicated psychiatric traits such as depressive symptoms, neuroticism, major depressive disorder and schizophrenia. These results point to a heritable component of suicide attempt and an incomplete genetic relationship with any single psychiatric disorder. In addition to identifying significant genetic correlations with previously implicated psychiatric traits we have identified two significant genetic correlations with non-psychiatric traits. The positive relationship between genetic risks of suicide and insomnia has been well studied with consistent evidence of the effect of disturbed sleep on suicidal behaviors including direct predictive effect of insomnia after accounting for depressive symptoms and other psychiatric traits^39,40^. Here, we demonstrate that these two traits share a common genetic risk profile pointing to shared underlying biology consistent with previous work showing genetic correlation of insomnia and other sleep traits and psychiatric disorders^34,41^. Further work will be needed to dissect potential independent genetic components across these related phenotypes. Additionally, we’ve identified an inverse relationship between the genetic risks of suicide and age at first birth. There is literature demonstrating increased risk of suicide among young mothers even after accounting for social status but this connection is not well-studied^42^. Age at first birth is also genetically and phenotypically correlated with many other traits making further interpretation of this result difficult.

We demonstrate both a polygenic risk signature and a genetic correlation between patient-reported suicide attempt and a clinically-predicted risk of attempting suicide. These results demonstrate that clinically predicted probability of attempting suicide based only on EHR data has a genetic component that is comparable to the patient-reported phenotype. Further, we have created a quantitative value for a binary trait that significantly correlates with a quantitative measure of genetic risk. The quantitative measure allows us to significantly increase our power for genetic studies demonstrated by comparable estimates of heritability between the 337,000 individual UK Biobank sample with 2,400 cases and the 24,000 individual BioVU sample with 73 validated cases. The opportunities to extend this approach to additional phenotypes that are hard to ascertain or rare are extensive but, of course, require the presence of substantial clinical data and enough validated cases to successfully predict those outcomes. While we had only 24,000 samples with the genetic data from BioVU we had access to 2.8M patients in the VUMC EHR including 3,250 chart confirmed cases of suicide attempt to perform prediction and calculate posterior probabilities. Clinical data are becoming more available, more accessible, more detailed and integrated into larger systems further enabling this approach to power future genetic studies of many phenotypes.

Despite the increase in power from using the posterior probabilities, we identified only two genome-wide significant loci, both of which did not replicate in the UK Biobank sample and no genome-wide significant loci were identified in the UK Biobank sample alone. For genetic studies of suicide, the samples sizes used here were large but these numbers are small when compared to the number of samples required before other psychiatric and non-psychiatric phenotypes successfully identified loci with GWAS. Based on the polygenic analysis and the significant estimates of h^2^SNP, we anticipate the identification of many genome-wide significant loci of suicidal behavior as sample sizes continue to increase. Further, while the genetic correlation between the two suicide attempt phenotypes is high the standard error is large leaving a substantial confidence interval ranging as low as 0.71. The differences seen in genetic correlations with intelligence and educational attainment may be a product of sample population or ascertainment which differ between a hospital population and a national population sample restricted to individuals between 40 and 69. Further work and increased sample sizes will be required to determine the true genetic correlation of these phenotypes and how they differ.

Taken together, our results point to a significant genetic component of suicide attempt and a significant but incomplete genetic relationship with psychiatric and sleep traits. We demonstrate that utilizing clinical data from EHR and machine learning approaches can generate quantitative risk probabilities that share substantial genetic etiology to a more classically used patient-reported outcome. Finally, we show that these quantitative probabilities can be used to substantially improve the power of genetic studies relative to relying on the binary trait alone.

## Author contributions

DMR designed and conceived the study. CGW and DMR generated the quantitative suicide attempt phenotype. DMR and MWA performed analyses. DMR, MAR, CGW, JDR and JCF provided interpretation of results. DMR drafted the manuscript. DMR, CGW, MAR, JDR and JCF provided critical revisions of the manuscript. All authors have read and approve submission.

## Acknowledgements

This research has been conducted using the UK Biobank resource. We thank the participants of this study. The primary and processed data used in these analyses are available in the UK Biobank access management system (https://amsportal.ukbiobank.ac.uk/) under application 24983, “Generating effective therapeutic hypotheses from genomic and hospital linkage data” (http://www.ukbiobank.ac.uk/wp-content/uploads/2017/06/24983-Dr-ManuelRivas.pdf). The dataset(s) used for the analyses described were obtained from Vanderbilt University Medical Center’s BioVU which is supported by numerous sources: institutional funding, private agencies, and federal grants. These include the NIH funded Shared Instrumentation Grant S10RR025141; and CTSA grants UL1TR002243, UL1TR000445, and UL1RR024975. Genomic data are also supported by investigator-led projects that include U01HG004798, R01NS032830, RC2GM092618, P50GM115305, U01HG006378, U19HL065962, R01HD074711; and additional funding sources listed at https://victr.vanderbilt.edu/pub/biovu/. Y.T. is supported by Funai Overseas Scholarship from Funai Foundation for Information Technology and the Stanford University Biomedical Informatics Training Program. We thank Dr. Nancy Cox, Dr. Eli Stahl, Dr. Alex Charney and Andrew Kirby for helpful discussion and comments on the manuscript.

## References

1. WHO | Suicide data. WHO Available at: http://www.who.int/mental_health/prevention/suicide/suicideprevent/en/. (Accessed: 3rd January 2018)

2. Statistics|Suicide|Violence Prevention|Injury Center|CDC. Available at: https://www.cdc.gov/violenceprevention/suicide/statistics/. (Accessed: 3rd January 2018)

3. Suicidal Thoughts and Behaviors Among Adults Aged ≥18 Years ‑‑‑ United States, 2008-2009. Available at: https://www.cdc.gov/mmwr/preview/mmwrhtml/ss6013a1.htm. (Accessed: 10th January 2018)

4. Nock, M. K. et al. Suicide and Suicidal Behavior. Epidemiol. Rev. 30, 133–154 (2008).

5. Olfson, M. et al. National Trends in Suicide Attempts Among Adults in the United States. JAMA Psychiatry 74, 1095–1103 (2017).

6. Statham, D. J. et al. Suicidal behaviour: an epidemiological and genetic study. Psychol. Med. 28, 839–855 (1998).

7. Roy, A. & Segal, N. L. Suicidal behavior in twins: a replication. J. Affect. Disord. 66, 71–74 (2001).

8. Sokolowski, M., Wasserman, J. & Wasserman, D. Genome-wide association studies of suicidal behaviors: A review. Eur. Neuropsychopharmacol. 24, 1567–1577 (2014).

9. Voracek, M. & Loibl, L. M. Genetics of suicide: a systematic review of twin studies. Wien. Klin. Wochenschr. 119, 463–475 (2007).

10. Perlis, R. H. et al. Genome-wide association study of suicide attempts in mood disorder patients. Am. J. Psychiatry 167, 1499–1507 (2010).

11. Stein, M. B. et al. Genomewide association studies of suicide attempts in US soldiers. Am. J. Med. Genet. B Neuropsychiatr. Genet. n/a-n/a doi:10.1002/ajmg.b.32594

12. Willour, V. L. et al. A genome-wide association study of attempted suicide. Mol. Psychiatry 17, 433 (2012).

13. Zai, C. C. et al. A genome-wide association study of suicide severity scores in bipolar disorder. J. Psychiatr. Res. 65, 23–29 (2015).

14. Chang, B. P. et al. Biological risk factors for suicidal behaviors: a meta-analysis. Transl. Psychiatry 6, e887 (2016).

15. Brent, D. A. et al. Psychiatric Risk Factors for Adolescent Suicide: A Case-Control Study. J. Am. Acad. Child Adolesc. Psychiatry 32, 521–529 (1993).

16. Mental disorders and comorbidity in suicide. Am. J. Psychiatry 150, 935–940 (1993).

17. Brent, D. A. & Mann, J. J. Family genetic studies, suicide, and suicidal behavior. Am. J. Med. Genet. C Semin. Med. Genet. 133C, 13–24 (2005).

18. Brent, D. A., Bridge, J., Johnson, B. A. & Connolly, J. Suicidal Behavior Runs in Families: A Controlled Family Study of Adolescent Suicide Victims. Arch. Gen. Psychiatry 53, 1145–1152 (1996).

19. Barak-Corren, Y. et al. Predicting Suicidal Behavior From Longitudinal Electronic Health Records. Am. J. Psychiatry 174, 154–162 (2016).

20. Walsh, C. G., Ribeiro, J. D. & Franklin, J. C. Predicting Risk of Suicide Attempts Over Time Through Machine Learning. Clin. Psychol. Sci. 216770261769156 (2017). doi:10.1177/2167702617691560

21. McCoy, T. H., Castro, V. M., Roberson, A. M., Snapper, L. A. & Perlis, R. H. Improving Prediction of Suicide and Accidental Death After Discharge From General Hospitals With Natural Language Processing. JAMA Psychiatry 73, 1064–1071 (2016).

22. Kessler, R. C. et al. Predicting suicides after outpatient mental health visits in the Army Study to Assess Risk and Resilience in Servicemembers (Army STARRS). Mol. Psychiatry (2016). doi:10.1038/mp.2016.110

23. Kessler, R. C. et al. Predicting Suicides After Psychiatric Hospitalization in US Army Soldiers: The Army Study to Assess Risk and Resilience in Servicemembers (Army STARRS). JAMA Psychiatry 72, 1–9 (2014).

24. Bycroft, C. et al. Genome-wide genetic data on ∼500,000 UK Biobank participants. bioRxiv 166298 (2017). doi:10.1101/166298

25. Roden, D. et al. Development of a Large-Scale De-Identified DNA Biobank to Enable Personalized Medicine. Clin. Pharmacol. Ther. 84, 362–369 (2008).

26. Harrell, F. E. J. Regression Modeling Strategies. Medicine (Baltimore) (2006). doi:10.1007/978-1-4757-3462-1

27. Smith, G. C. S., Seaman, S. R., Wood, A. M., Royston, P. & White, I. R. Correcting for optimistic prediction in small data sets. Am. J. Epidemiol. 180, 318–324 (2014).

28. Bulik-Sullivan, B. K. et al. LD Score regression distinguishes confounding from polygenicity in genome-wide association studies. Nat. Genet. 47, 291–295 (2015).

29. Purcell, S. et al. PLINK: A Tool Set for Whole-Genome Association and Population-Based Linkage Analyses. Am. J. Hum. Genet. 81, 559–575 (2007).

30. Zheng, J. et al. LD Hub: a centralized database and web interface to perform LD score regression that maximizes the potential of summary level GWAS data for SNP heritability and genetic correlation analysis. Bioinformatics 33, 272–279 (2017).

31. Okbay, A. et al. Genetic variants associated with subjective well-being, depressive symptoms, and neuroticism identified through genome-wide analyses. Nat. Genet. **advance online publication,** (2016).

32. Schizophrenia Working Group of the Psychiatric Genomics Consortium. Biological insights from 108 schizophrenia-associated genetic loci. Nature 511, 421–427 (2014).

33. Hammerschlag, A. R. et al. Genome-wide association analysis of insomnia complaints identifies risk genes and genetic overlap with psychiatric and metabolic traits. Nat. Genet. **advance online publication,** (2017).

34. Lane, J. M. et al. Genome-wide association analyses of sleep disturbance traits identify new loci and highlight shared genetics with neuropsychiatric and metabolic traits. Nat. Genet. 49, 274 (2017).

35. Consortium, M. D. D. W. G. of the P. G. et al. A mega-analysis of genome-wide association studies for major depressive disorder. Mol. Psychiatry 18, 497 (2013).

36. Consortium, C.-D. G. of the P. G. Identification of risk loci with shared effects on five major psychiatric disorders: a genome-wide analysis. The Lancet 381, 1371–1379 (2013).

37. Barban, N. et al. Genome-wide analysis identifies 12 loci influencing human reproductive behavior. Nat. Genet. 48, 1462 (2016).

38. Okbay, A. et al. Genome-wide association study identifies 74 loci associated with educational attainment. Nature 533, 539–542 (2016).

39. Pigeon, W. R., Pinquart, M. & Conner, K. Meta-Analysis of Sleep Disturbance and Suicidal Thoughts and Behaviors. J. Clin. Psychiatry 73, 1160–1167 (2012).

40. Ribeiro, J. D. et al. Sleep problems outperform depression and hopelessness as cross-sectional and longitudinal predictors of suicidal ideation and behavior in young adults in the military. J. Affect. Disord. 136, 743–750 (2012).

41. Hammerschlag, A. R. et al. Genome-wide association analysis of insomnia complaints identifies risk genes and genetic overlap with psychiatric and metabolic traits. Nat. Genet. **advance online publication,** (2017).

42. Otterblad Olausson, P., Haglund, B., Ringbäck Weitoft, G. & Cnattingius, S. Premature death among teenage mothers. BJOG Int. J. Obstet. Gynaecol. 111, 793–799 (2004).

